# Multi-scale approaches for high-speed imaging and analysis of large neural populations

**DOI:** 10.1101/091132

**Authors:** Johannes Friedrich, Weijian Yang, Daniel Soudry, Yu Mu, Misha B. Ahrens, Rafael Yuste, Darcy S. Peterka, Liam Paninski

## Abstract

Progress in modern neuroscience critically depends on our ability to observe the activity of large neuronal populations with cellular spatial and high temporal resolution. However, two bottlenecks constrain efforts towards fast imaging of large populations. First, the resulting large video data is challenging to analyze. Second, there is an explicit tradeoff between imaging speed, signal-to-noise, and field of view: with current recording technology we cannot image very large neuronal populations with simultaneously high spatial and temporal resolution.

Here we describe multi-scale approaches for alleviating both of these bottlenecks. First, we show that spatial and temporal decimation techniques provide order-of-magnitude speedups in spatiotemporally demixing calcium video data into estimates of single-cell neural activity. Second, once the shapes of individual neurons have been identified (e.g., after an initial phase of conventional imaging with standard temporal and spatial resolution), we find that the spatial/temporal resolution tradeoff shifts dramatically: after demixing we can accurately recover neural activity from data that has been spatially decimated by an order of magnitude. This offers a cheap method for compressing this large video data, and also implies that it is possible to either speed up imaging significantly, or to “zoom out” by a corresponding factor to image order-of-magnitude larger neuronal populations with minimal loss in accuracy or temporal resolution.

**Author Summary:** The voxel rate of imaging systems ultimately sets the limit on the speed of data acquisition. These limits often mean that only a small fraction of the activity of large neuronal populations can be observed at high spatio-temporal resolution. For imaging of very large populations with single cell resolution, temporal resolution is typically sacrificed. Here we propose a multi-scale approach to achieve single cell precision using fast imaging at reduced spatial resolution. In the first phase the spatial location and shape of each neuron is obtained at standard spatial resolution; in the second phase imaging is performed at much lower spatial resolution. We show that we can apply a demixing algorithm to accurately recover each neuron’s activity from the low-resolution data by exploiting the high-resolution cellular maps estimated in the first imaging phase. Thus by decreasing the spatial resolution in the second phase, we can compress the video data significantly, and potentially acquire images over an order-of-magnitude larger area, or image at significantly higher temporal resolution, with minimal loss in accuracy of the recovered neuronal activity. We evaluate this approach on real data from light-sheet and 2-photon calcium imaging.

## Introduction

A major goal of neuroscience is to understand interactions within large populations of neurons, including their network dynamics and emergent behavior. This ideally requires the observation of neural activity over large volumes. Recently, light-sheet microscopy and genetically encoded indicators have enabled unprecedented whole-brain imaging of tens of thousands of neurons at cellular resolution [1]. However, light-sheet microscopy generally suffers from slow volumetric speeds (e.g. [2], but see also [3,4]) and is usually applied to small and transparent brains. In scattering brains, current technologies with single-neuron resolution are usually based on slow, serially-scanned two-photon (2P) imaging methods that can only sample from *O*(10^2^ – 10^3^) neurons simultaneously with adequate temporal resolution [5]. Recent advances have enabled faster light-sheet imaging in cortex [6] and fast volumetric 2P imaging [7], but we must still contend with critical trade-offs between temporal and spatial resolution — and the need for even faster imaging of even larger neural populations.

Another critical challenge is the sheer amount of data generated by these large-scale imaging methods. A crucial step for further neural analysis involves a transition from voxel-space to neuron-source space: i.e., we must detect the neurons and extract and demix each neuron’s temporal activity from the video. Simple methods such as averaging voxels over distinct regions of interest (ROIs) are fast, but more statistically-principled methods based on constrained non-negative matrix factorization (CNMF) better conserve information, yield higher signal-to-noise ratio, recover more neurons, and enable the demixing of spatially overlapping neurons [8]. The methods described in [8] were not optimized for very large datasets, but NMF is a key machine learning primitive that has enjoyed more than a decade of intensive algorithmic optimization [9–12] that we can exploit here to scale the CNMF approach. We find that a very simple idea leads to order-of-magnitude speedups: by decimating the data (i.e., decreasing the resolution of the data by simple local averaging [13]), we can obtain much faster algorithms with minimal loss of accuracy.

Decimation ideas do not just lead to faster computational image processing, but also offer prescriptions for faster image acquisition over larger fields of view (FOV), and for observing larger neural populations. Specifically, we propose the following two-phase combined image acquisition/analysis approach. In the first phase, we use conventional imaging methods to obtain estimates of the visible neuronal locations and shapes. After this cell-identification phase is complete we switch to low-spatial-resolution imaging, which in the case of camera-based imaging simply corresponds to “zooming out” on the image, i.e., expanding the spatial size of each voxel. This has the benefit of projecting a larger FOV onto the same number of voxels; alternatively, if the number of voxels recorded per second is a limiting factor, then recording fewer (larger) voxels per frame implies that we can image at higher frame-rates. We are thus effectively trading off spatial resolution for temporal resolution; if we cut the spatial resolution too much we may no longer be able to clearly identify or resolve single cells by eye in the obtained images. However, we show that, given the high-spatial-resolution information obtained in the first imaging phase, the demixing stage of CNMF can recover the temporal signals of interest even from images that have undergone radical spatial decimation (an order of magnitude or more). In other words, CNMF significantly shifts the tradeoff between spatial and temporal resolution, enabling us to image larger neuronal populations at higher temporal resolution.

The rest of this paper is organized as follows. We first describe how temporal and spatial decimation (along with several other improvements) can be used within the CNMF algorithm to gain order-of-magnitude speed-ups in calcium imaging video processing. Next we investigate how decimation can enable faster imaging of larger populations for light-sheet and 2P imaging. We show the importance of the initial cell identification phase, quantitatively illustrate how CNMF changes the tradeoff between spatial and temporal resolution, and discuss how spatially decimated imaging followed by demixing can be interpreted as a simple compression and decoding scheme. We show that good estimates of the neural shapes can be obtained on a small batch of standard-resolution data, corresponding to a short cell-identification imaging phase. Finally we demonstrate that interleaved imaging that translates the pixels by subpixel shifts on each frame further improves the fidelity of the recovered neural time series.

## Results

### Order-of-magnitude speedups in demixing calcium imaging data

Constrained non-negative matrix factorization (see Methods) relies on the observation that the spatiotemporal fluorescence activity (represented as a space-by-time matrix) can be expressed in terms of a product of two matrices: a spatial matrix *A* that encodes the location and shape of each neuron and a temporal matrix *C* that characterizes the calcium concentration within each neuron over time. Placing constraints on the spatial footprint of each neuron (e.g., enforcing sparsity and locality of each neural shape) and on the temporal activity (modeling the observed calcium in terms of a filtered version of sparse, non-negative neural activity) significantly improves the estimation of these components compared to vanilla NMF [8]. Below we describe a number of algorithmic improvements on the basic approach described in [8]: an iterative block-coordinate descent algorithm in which we optimize for components of *A* with *C* held fixed, then for *C* with *A* held fixed.

We begin by considering imaging data obtained at low temporal resolution, specifically a whole-brain light-sheet imaging recording acquired at a rate of 2 Hz using nuclear localized GCaMP6f in zebrafish. We restricted our analysis to a representative patch shown in Fig 2A, extracted from a medial z-layer of the telencephalon (pallium). (Similar analyses were also performed on patches from midbrain and hindbrain, with similar conclusions.) The neural centers were detected automatically using the greedy method from [8]. To ensure that the spatial components in *A* are localized, we constrained them to lie within spatial sub-patches (dashed squares in Fig 2A; see also Methods).

The first algorithmic improvement follows from the realization that some of the constraints applied in CNMF are unnecessary, at least during early iterations of the algorithm, when only crude estimates for *A* and *C* are available. Specifically, [8] imposed temporal constraints on *C* in each iteration: namely, *C* was modeled as a filtered version of a nonnegative neural activity signal *S* — i.e., *CG = S*, for an invertible matrix *G* — and therefore *CG* is constrained to be non-negative. We found that enforcing a simpler non-negativity constraint on *C* instead of *CG* (and then switching to impose the constraint on *CG* only once the estimates of *A* and *C* were closer to convergence) led to a simpler algorithm enabling faster early iterations with no loss in accuracy (data not shown).

Next we found that significant additional speed-ups in this simplified problem could be obtained by simply changing the order in which the variables in this simplified block-coordinate descent scheme are updated [12]. Instead of updating the temporal activity and spatial shape of one neuron at a time (Fig 1A, black line) as in [8], which is known as hierarchical alternating least squares (HALS, [9]) or rank-one residue iteration (RRI, [14]), it turned out to be beneficial to update the activities of all neurons while keeping their shapes fixed, and then updating all shapes while keeping their activities fixed (Fig 1A, vermilion line). The ensuing method is a constrained version of the fast hierarchical alternating least squares (fast HALS, [15]) for NMF; one major advantage of this update ordering is that in each iteration we operate on smaller matrices obtained as dot-products of the data matrix *Y* with *A* or *C*, and there is no need to compute the large residual matrix *Y* – *AC* (which is of the same size as the original video) [10,11]. (In the comparisons below we computed the residual to quantify performance, but excluded the substantial time spent on its computation from the reported wall time values.)

**Fig 1.**
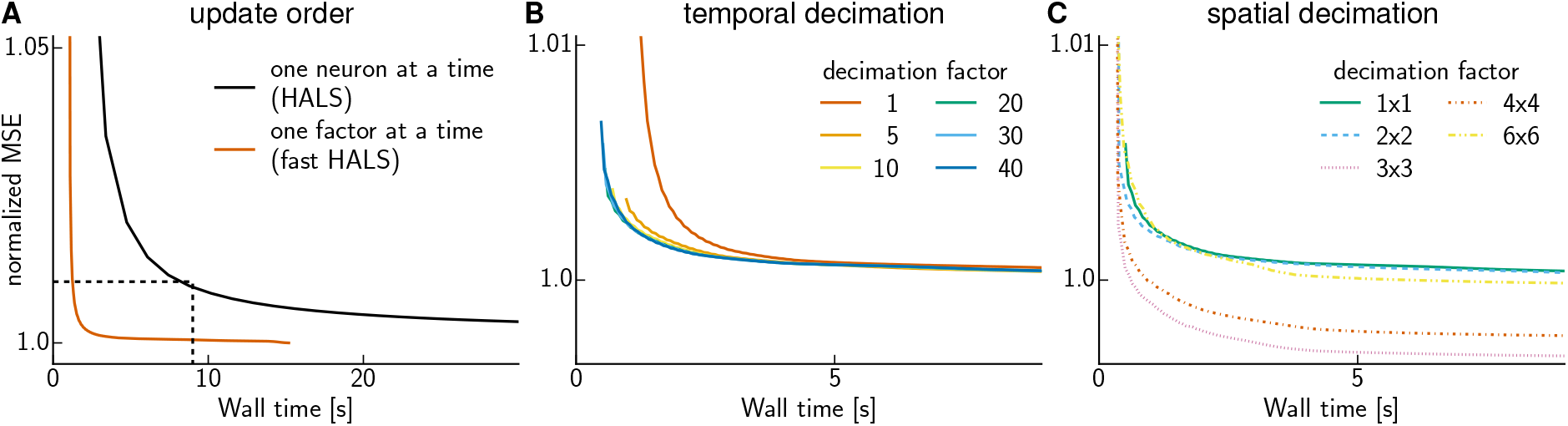
Speeding up CNMF. Mean-squared-error as a function of wall time. **(A)** The order in which coordinates in the block-coordinate descent method are updated affects convergence speed. The dashed box indicates the zoomed in plot region of (B,C) **(B)** Temporal decimation speeds up CNMF. Curves show the normalized mean square error (MSE; normalized so that at convergence the original CNMF method reaches a value of 1) on the whole data after 30 initial iterations are performed on a smaller dataset obtained by temporally decimating by different factors *k* — i.e., averaging each block of *k* frames into a single frame (color indicates *k*, cf. legend). The curve for decimation factor 1 (i.e., no decimation) is identical to the one in (A). **(C)** Spatial decimation (locally averaging over spatial blocks of size *l* × *l* in the image) yields further improvements, in particular a better (local) minimum of the MSE. Curves show the results for 30 initial iterations on data that has been decimated temporally by a factor of 30 and spatially by the factors *l* given in the legend. The solid green curve for spatial decimation factor 1 × 1 (i.e., no spatial decimation) is the same as in (B).

Next we reasoned that to obtain a good preliminary estimate of the spatial shape matrix *A*, it is likely unnecessary to use the original data at full temporal resolution [13]. Thus we experimented with the following approach: downsample temporally by a factor of *k*, then run constrained fast HALS (as described above) for 30 iterations, and then finally return to the original (non-downsampled) data and run a few more iterations of fast HALS until convergence. We experimented with three different downsampling methods: 1) selection of the *k*-th frame (this could be considered a kind of stochastic update rule, since we are forming updates based only on a subset of the data); 2) forming a median over the data in each block of *k* frames (applying the median over each pixel independently); and 3) forming a mean over each block of *k* frames. The mean approach (3) led to significantly more accurate and stable results than did the subsampling approach (1), consistent with the results of [12], and was about an order of magnitude faster than the median approach (2) with similar accuracy, so we restrict our attention to the mean approach (3) for the remainder of this work. (A further advantage of approach (3) relative to (1) is that (1) can miss fast activity transients.) Fig 1B shows the results obtained for a varying number of decimation factors *k*; we conclude that temporal decimation provides another significant speedup over the results shown in Fig 1A.

Further speed gains were obtained when applying spatial decimation (computing a mean within *l* × *l* pixel blocks) in addition to temporal decimation over the 30 preliminary fast HALS iterations (Fig 1C); see Algorithm 1 for full details. Strikingly, spatial decimation led not only to faster but also to better solutions (where solution quality is measured by the residual sum of square errors, ‖*Y* − *AC*‖^2^), apparently because the spatially-decimated solutions are near better local optima in the squared-error objective function than are the non-decimated solutions.

#### Algorithm 1 NMF algorithm using localization constraints and spatio-temporal decimation

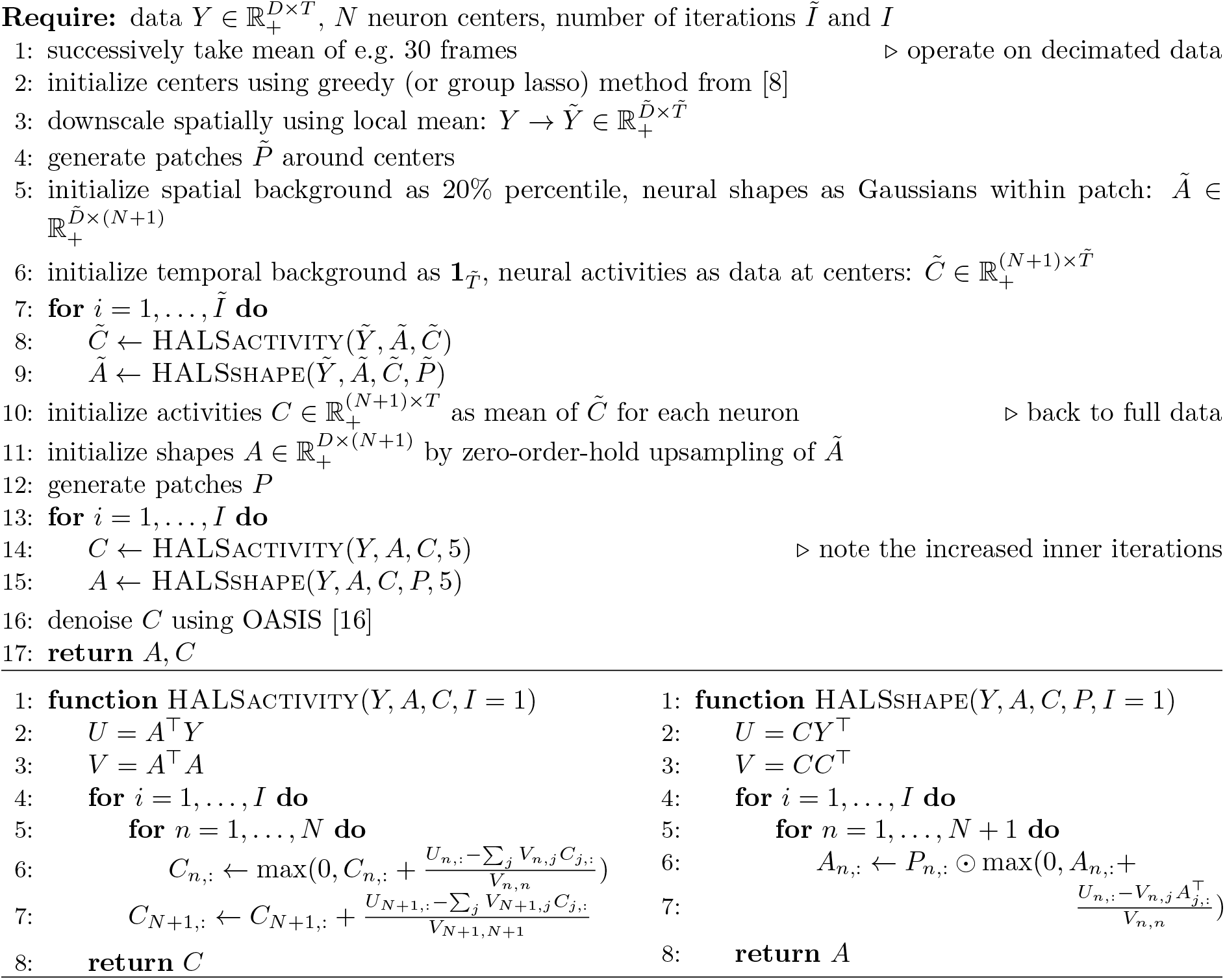

In summary, by simplifying the early iterations of the CNMF algorithm (by removing the temporal deconvolution constraints to use fast HALS iterations on temporally and spatially subsampled data), we obtained remarkable speed-ups without compromising the accuracy of the obtained solution, at least in terms of the sum-of-squares objective function. But how do these modifications affect the extracted neural shapes and activity traces? We ran the algorithm without decimation until convergence and with decimation for 1 and 10 s respectively. Fig 2 shows the results for three neurons with overlapping patches. Both shapes and activity traces agree well even if the decimated algorithm is run for merely 1 s (Fig 2B,C) and are nearly identical if run longer (Fig 2D); hence, decimation does not impair the final obtained accuracy.

**Fig 2.**
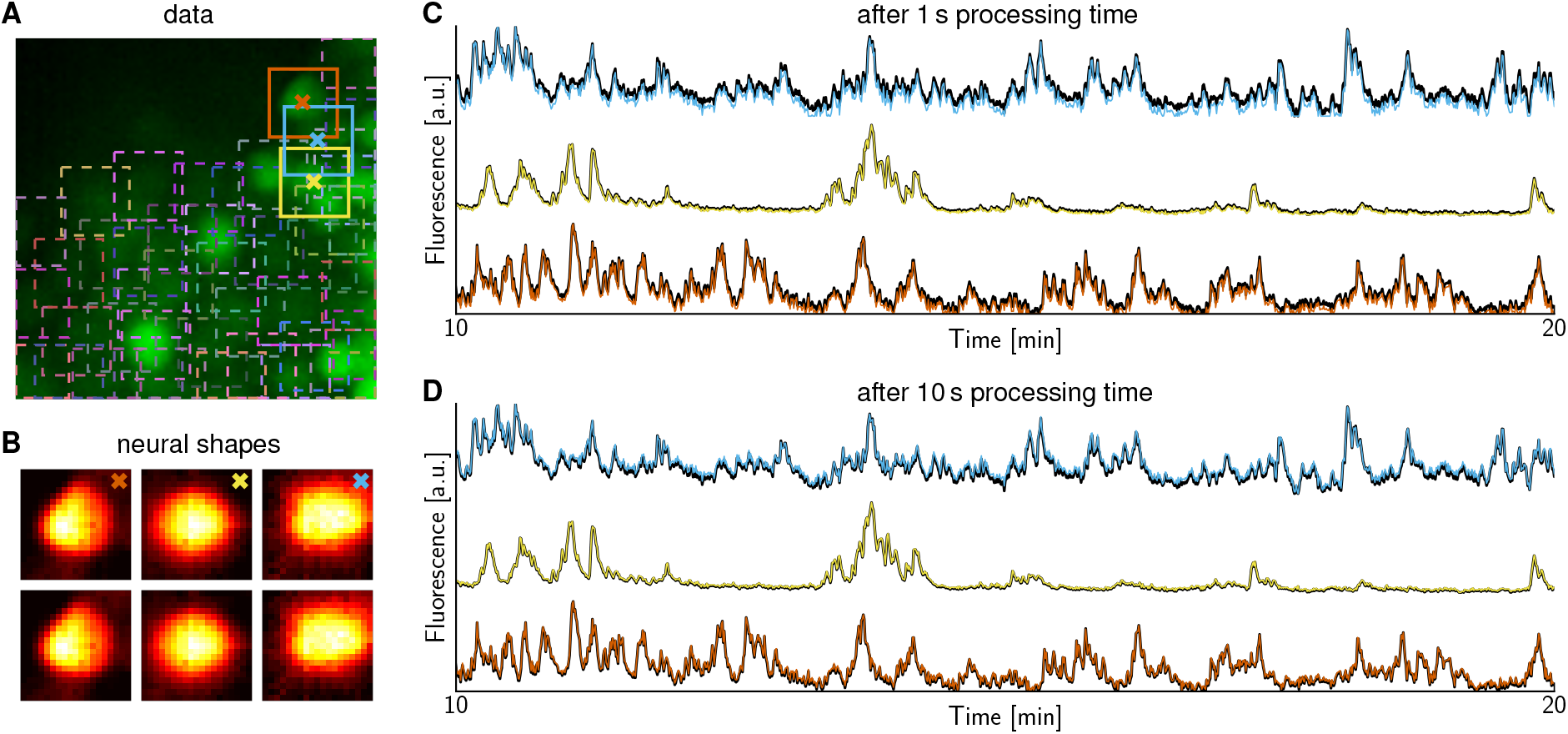
CNMF with and without initial decimation to speed up CNMF. **(A)** Real data summary image obtained as max-projection along time-axis. Squares indicate patches centered at suspected neurons. The three highlighted neurons are considered in the next panels. **(B)** Extracted shapes of the three neurons highlighted in (A) using the same color code. The upper row shows the result with initial temporal decimation by a factor of 30 and spatial decimation by 3×3, followed by five final iterations on the whole data. The algorithm ran for merely 1 s. The lower row shows the result without any decimation and running until convergence, yielding virtually identical results. **(C)** Extracted time traces without (thick black) and with initial iterations on decimated data (color as in A) overlap well after merely 1 s. **(D)** They overlap almost perfectly if the algorithm using decimation is run not only for 1 but 10 s.

Our focus has been on speeding up CNMF, one computational bottleneck of the entire processing pipeline. For completeness, we report the times spent on each step of the pipeline in Table 1 and compare to the previous CNMF version of [8]. After loading, the data was decimated temporally by a factor of 30 to speed up the detection of the neural centers using the greedy initialization method from [8]. We further decimated spatially and ran fast HALS for 30 iterations before finally returning to the whole data and performing five final fast HALS iterations. Each trace was normalized by the fluorescence at the resting state (known as Δ*F/F*) to account for baseline drift using a running percentile filter. Finally, the fluorescence traces were denoised via sparse non-negative deconvolution, using the recently developed fast method of [16], which eliminated another computational bottleneck present in the original CNMF implementation (last row of Table 1).

**Table 1:** Timing of the entire pipeline for a zebrafish patch.

| CNMF method | [8] | here & [16] |
| --- | --- | --- |
| load data<sup>†</sup> | 22 ± 1 | 22 ± 1 |
| decimate | N/A | 33 ± 1 |
| detect ROIs<sup>‡</sup> | 5,360 ± 20 | 135 ± 2 |
| NMF | 35,100 ± 400 | 990 ± 10 |
| $\Delta F/F$ | 410 ± 10 | 410 ± 10 |
| denoise | 17,900 ± 300 | 600 ± 20 |

Average computing time (± SEM) in ms over ten runs for individual steps of the processing pipeline on a standard laptop. The 96×96 patch contained *N* = 46 neurons recorded for *T* = 3000 frames. ^†^Loading the whole data as single binary file; loading a frame at a time was an order of magnitude slower. ^‡^Using greedy initialization; group lasso initialization [8] was an order of magnitude slower.

### Decimation can enable faster imaging of larger populations

We have shown that decimation leads to much faster computational processing of calcium video data. More importantly, these results inspired us next to propose a method for faster image acquisition or for imaging larger neural populations. The basic idea is quite simple: if we can estimate the quantities of interest (*A* and *C*) well given decimated data, then why collect data at the full resolution at all? Since spatial decimation by a factor of *l* conceptually reduces the number of pixels recorded over a given FOV by a factor of *l*^2^ (though of course this situation is slightly more complex in the case of scanning two-photon imaging; we will come back to this issue below), we should be able to use our newly-expanded pixel budget to image more cells, or image the same population of cells faster.

As we will see below, this basic idea can be improved upon significantly: if we have a good estimate for the spatial neural shape matrix *A* at the original spatial resolution, then we can decimate more drastically (thus increasing this *l*^2^ factor) with minimal loss in accuracy of the estimated activity *C*. This, finally, leads to the major proposal of this paper: first perform imaging with standard spatial resolution via conventional imaging protocols. Next perform the ROI detection and CNMF described above to obtain a good estimate of *A*. Then begin acquiring spatially *l*-decimated images and use the *C*-estimation step of CNMF to extract and demix the imaged activity. As we will see below, this two-phase imaging approach can potentially enable the accurate extraction of demixed neural activity even given quite large decimation factors *l*, with a correspondingly large increase in the resulting “imaging budget.”

#### Light-sheet imaging

We began by quantifying the potential effectiveness of this strategy using the zebrafish light-sheet imaging data examined in the last section. We emulated the decrease in spatial resolution by decimating the original imaging data as well as the neural shapes we had obtained by the CNMF approach, with a variety of decimation factors *l*. Then we used these decimated shapes *A*_*l*_ to extract and demix the activities *C*_*l*_ from the corresponding downscaled data *Y*_*l*_. We evaluated the resulting *C*_*l*_ traces by comparing them to the original traces *C*_1_ obtained from the full original data *Y*. (In our notation if *l =* 1 then no decimation is applied.) Fig 3A shows that similar fluorescence traces are inferred even from quite heavily coarsened shapes *A*_*l*_. The correlation between *C*_1_ and *C*_*l*_ decreases gracefully with the decimation factor *l* (Fig 3B, orange line). In contrast, this correlation drops precipitously for relatively small values of *l* in the ‘single-phase’ imaging setting (cyan line), where we estimate the shapes *A*_*l*_ directly from *Y*_*l*_, instead of estimating *A*_1_ from the full data *Y* first and then decimating to obtain *A*_*l*_. The problem in this ‘single-phase’ imaging setting is that ROI detection fails catastrophically once the pixelization becomes too coarse.

**Fig 3.**
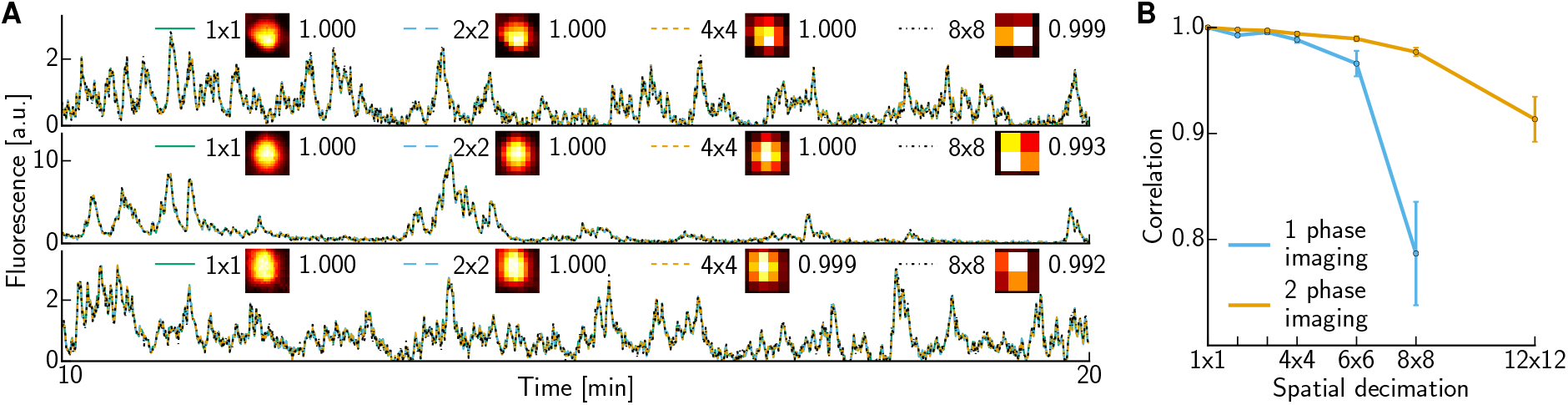
Low spatial resolution light-sheet imaging with previously identified cells. (A) Inferred traces for the three neurons in Fig 2. Shapes were decimated by averaging 2×2, 4×4 or 8×8 pixels. The legend shows the resulting shapes *A*_*l*_ as well as the correlation between the calcium traces *C*_*l*_ inferred from the decimated data versus the original traces *C*_1_ obtained from the non-decimated data. **(B)** Correlation between *C*_*l*_ and *C*_*l*_ for all cells recovered from this patch. The average correlation (±SEM) decays slowly as *l* increases for 2-phase imaging (orange) but drops abruptly if the shapes *A*_*l*_ are estimated directly from decimated data (cyan) instead of being decimated from the shapes *A*_1_ estimated from a standard-resolution imaging phase.

Thus the results are quite promising; with the two-phase imaging strategy we can effectively increase our imaging budget (as measured by *l*^2^) by over an order of magnitude with minimal loss in the accuracy of the obtained activity traces *C*_*l*_, and we also observe clear advantages of the two-phase over the single-phase decimation approaches. Finally, S1 Video illustrates the results in video form; we see there that we can recover essentially all the relevant information (at least visually) in the original video data *Y* from the spatially decimated video *Y*_*l*_, even with quite large decimation levels *l* (*l =* 8 in S1 Video). Thus this decimation-then-demix approach could also provide a trivial compression scheme (with the estimation of *C*_*l*_ from *Y*_*l*_ and *A*_*l*_ serving as the decoder; here the compression ratio is *l*^2^) that could be useful e.g. for wireless recordings, or any bandwidth- or memory-limited pipeline [17–19].

#### Two-photon imaging

Turning towards imaging data acquired at a faster frame-rate, we next consider the case of a 2P calcium imaging dataset from mouse visual cortex acquired at 20 Hz. We chose ROIs based on the correlation image and max-projection image, cf. Fig 4A and S2 Video, and then obtained neural shapes *A*_1_ and fluorescence activity *C*_1_ using CNMF. The upper row in Fig 4 (and S2 Video) shows the raw data *Y* and its reconstruction *A*_1_ · *C*_1_ based on CNMF, illustrating that all relevant ROIs have been detected and the matrix decomposition computed by CNMF captures the data well. The lower row shows the spatially decimated data *Y*_*l*_ and the reconstruction *A*_1_ · *C*_*l*_ based on sparse demixing of *Y*_*l*_ using knowledge of the neural shapes *A*_1_ from an initial cell identification imaging phase. As in the example in the last section, this illustrates that sparse demixing paired with cell shape identification applied to decimated data (panel D) captures the data virtually as well as CNMF without decimation (panel B). See S2 Video for full details.

**Fig 4.**
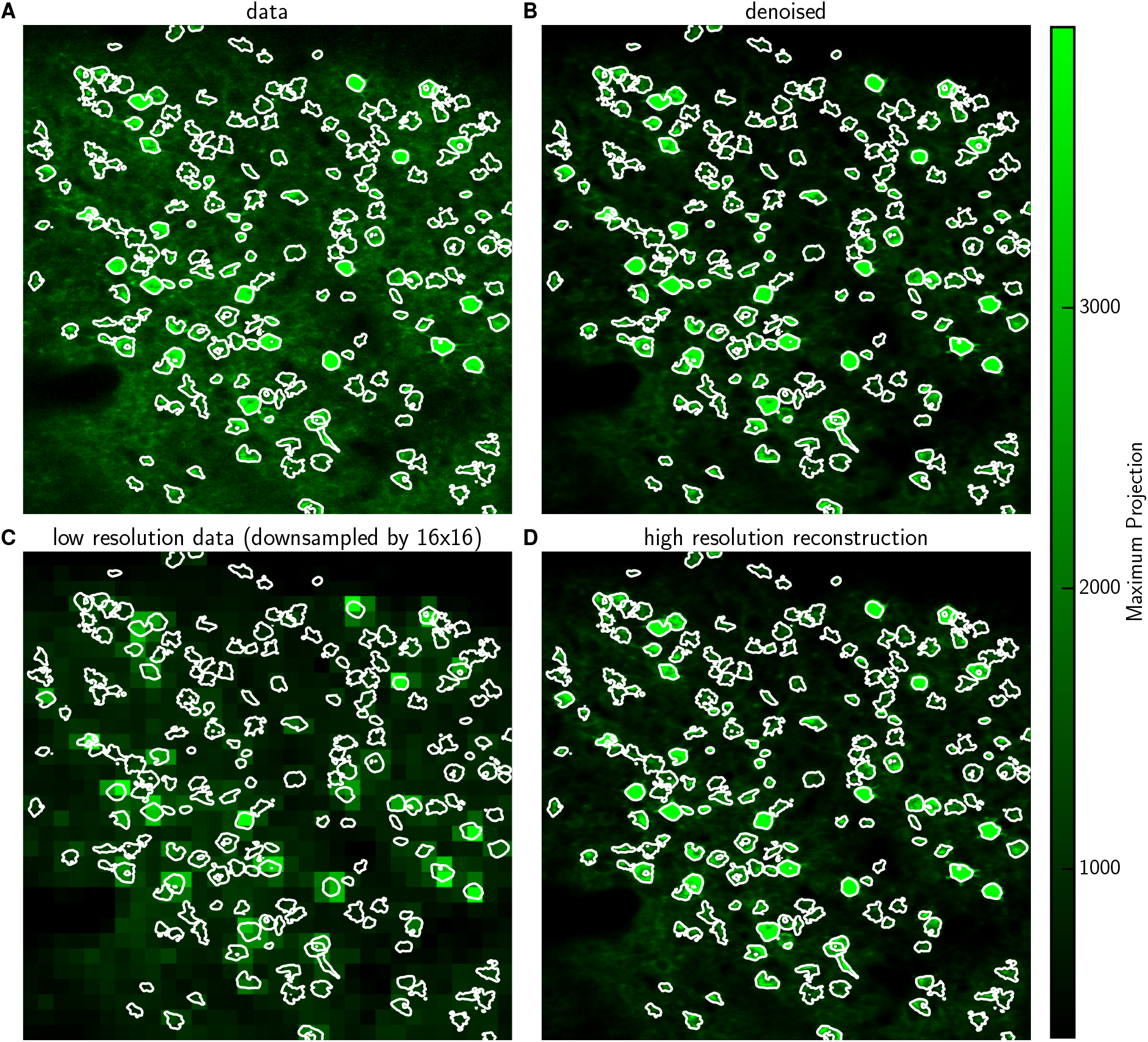
Previously identified shapes allow reconstruction at high spatial resolution based on low resolution imaging (best seen in S2 Video for full details). **(A)** Max projection image of the raw data *Y* with identified ROIs, i.e. neurons or activity hotspots. Contour lines contain 90% of the energy of each neural shape. **(B)** Max projection image of the denoised estimate *A*_1_ · *C*_1_ (plus the estimated background). **(C)** Max projection image for data obtained at lower spatial resolution, *Y*_*l*_; *l* = 16 here. **(D)** Reconstruction based on the low resolution data in (C) and previously identified shapes, *A*_1_ · *C*_*l*_. The reconstruction looks very similar to the denoised high-resolution data of (B). Note: contours in (B-D) are not recomputed in each panel, but rather are copied from (A), to aid comparison.

Fig 5 shows, analogously to Fig 3A, the traces *C*_*l*_ recovered from spatially decimated data (using the 2-phase imaging approach) depend quite weakly on the decimation factor *l*, until *l* becomes so large that the resulting pixel size is comparable to the size of individual somas. We also deconvolved the estimated calcium transients *C*_*l*_ into estimates of neural activity *S*_*l*_, using the sparse non-negative deconvolution method described in [8, 16]; we see similarly weak dependence of the results on *l* when comparisons are performed on this deconvolved activity. The ROIs were rank-sorted using a combination of maximal fluorescence intensity and a measure of the compactness of the neural shape (see Methods); we find that the similarity between *C*_1_ and *C*_*l*_ is strongest for the highest-ranked ROIs.

**Fig 5.**
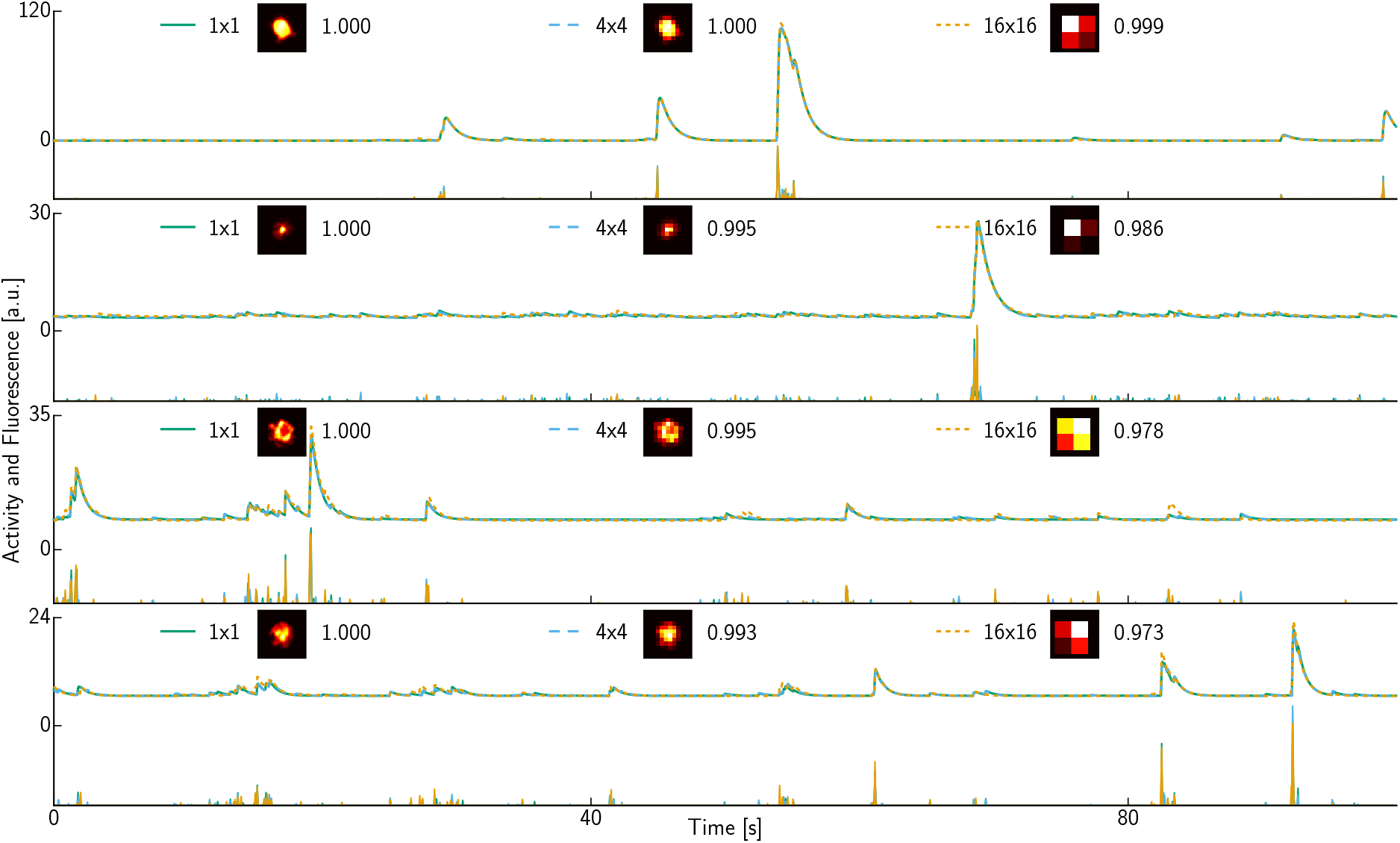
**Inferred denoised and deconvolved traces** for ROIs with rank 1, 40, 80 and 120 (top to bottom) out of a total of *N* = 187 ROIs in the 2P dataset. In each row, a denoised trace from *C*_*l*_ is shown above the corresponding deconvolved trace from *S*_*l*_. Shapes were decimated by averaging 4×4 or 16×16 pixels. The legend shows the resulting shapes *A*_*l*_ as well as the correlation of the inferred denoised fluorescence *C*_*l*_ versus the estimate *C*_1_ obtained without decimation.

We summarize the results over all neurons in this dataset in Fig 6. As in the examples shown in Fig 5, we see that the mean correlation between *C*_1_ and *C*_*l*_ decreases gracefully with *l* in the 2-phase imaging setting (orange lines). In contrast, the correlation decays much more sharply if the shapes *A*_*l*_ were obtained directly on low-resolution data (1 phase imaging; cyan lines), similarly but more dramatically than in Fig 3B. Similar results hold for the deconvolved activity *S*_1_ and *S*_*l*_ (dashed), though the correlation between *S*_1_ and *S*_*l*_ does decay more quickly than does the correlation between *C*_1_ and *C*_*l*_. Fig 6C shows that the decrease in correlation is more pronounced for lower ranked ROIs; i.e., if we restrict attention to the most clearly-identified cells then we can safely spatially decimate even more aggressively.

**Fig 6.**
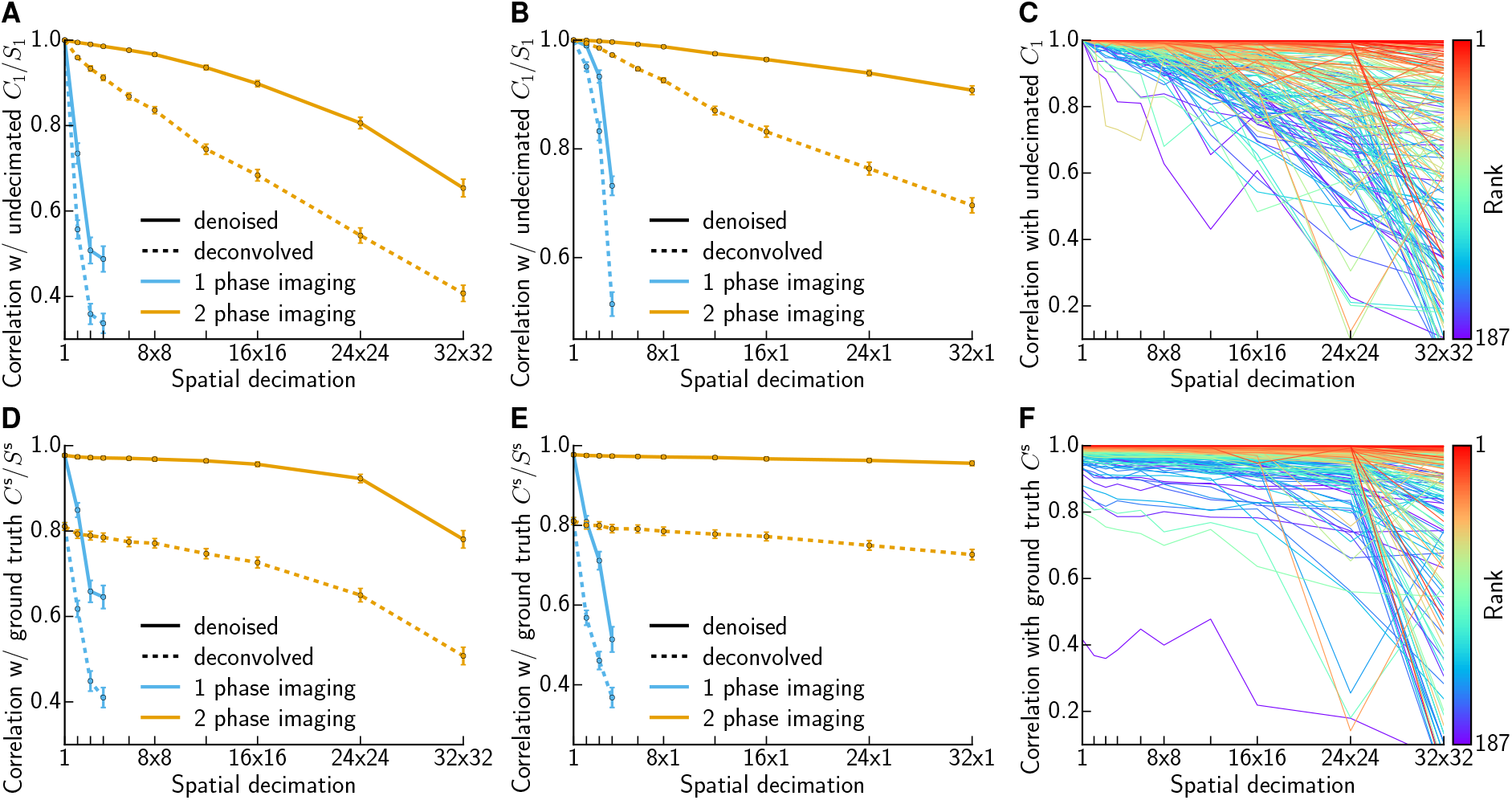
Quantifying the impact of decimation for 2P data. **(A)** Summary of correlations between denoised traces *C*_1_ and *C*_*l*_ and deconvolved traces *S*_1_ and *S*_*l*_. Decimating in *x* and *y* direction. The average correlation (±SEM) of denoised fluorescence (solid) decays slowly for 2 phase imaging (orange) and abruptly if the shapes *A*_*l*_ are not inferred in the pre-screening phase but are instead estimated directly from downscaled data (cyan). The same holds for the deconvolved traces (dashed). **(B)** Analogous to (A), but with decimation applied just in the spatial horizontal direction. **(C)** Correlation of denoised fluorescence for each ROI plotted individually. Rank of ROI indicated by color; better ranked ROIs are less susceptible to decimation errors. **(D-F)** Analogous to (A-C), but comparing to simulated ground truth *C*^s^ instead of inferred 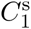. Traces were obtained on decimated simulated data with reshuffled residuals.

So far we have used the correlation between *C*_1_ and *C*_*l*_ (or *S*_1_ and *S*_*l*_) to quantify the robustness of signal recovery from the decimated data *Y*_*l*_. However, *C*_1_ and *S*_1_ should not be considered “ground truth”: these are merely estimates of the true underlying neural activity, inferred from noisy data *Y*, and it is not surprising that *C*_1_ and *C*_*l*_ are close for small values of *l*. A more critical question is the following: is *C*_*l*_ a significantly worse estimate of the true underlying neural activity *C* than *C*_1_? Of course ground truth neural activity is not available for this dataset, but we can simulate data *Y*^s^ and compare how well 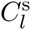 and 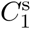 recover the simulated ground truth *C*^s^. To generate this simulated dataset, we started with *A*^s^:= *A*_1_ and *C*^s^:= *C*_1_ recovered from the full-resolution original data *Y*, and then generated a new simulated matrix *Y*^s^ = *A*^s^*C*^s^ + *B* + *R*^s^, where *B* represents background terms and *R*^s^ is chosen to match the statistics of the original residual *R* = *Y* − *A*_1_*C*_1_ − *B* (specifically, for each pixel *d* we formed 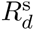 by randomly re-ordering the original corresponding residual time series *R*_*d*_ in that pixel; see Methods). Then we estimated 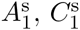 and 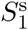 from *Y*^s^, and 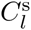 and 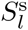 from decimated versions of *Y*^s^, and compared the results in Fig 6D-F. We see some important differences compared to Fig 6A-C: the correlation curves no longer approach 1 as the decimation level *l* decreases towards 1 (for example, the mean correlation between ground truth and recovered *S*^s^ is about 0.8 even if no decimation is applied), and the correlation between ground truth and the recovered 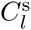 (or 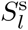) now decreases much more slowly as a function of *l —* indicating that signal recovery from spatially decimated data is even more robust than indicated in Fig 6A-C. On the other hand, we continue to observe similar strong differences in recovery when comparing the 1- and 2-phase imaging results.

In practice we envision that the first phase of cell identification will only be performed on a small initial batch of the data (and maybe also at the end of the experiment to check for consistency). Therefore we experimented with reducing the amount of data used to infer the neural shapes from the full data (2,000 frames, 100 s) to 1,000 frames (50 s) and 500 frames (25 s) respectively. Fig 7A shows the shapes inferred based on the full data (orange) or using only the first half (blue) with a max projection of the first half of the data as background. While some neural shapes are less well defined using less data, we only seem to miss two suspects entirely (white arrows). However, closer inspection reveals that these ROIs are not missed neurons, but rather are highly correlated with and actually part of the neuron indicated by the blue arrow — so in this example 50 seconds was sufficient to identify all the necessary neural shapes. Fig 7B,C report the correlation values measured only on the second half of the data, using shapes *A* estimated just with the first half. The close proximity of the orange and blue trace is further evidence that the shapes *A* can be estimated sufficiently in a short pre-screening phase. Interestingly, estimating the shapes based on merely 500 frames leads to some slight overfitting; the correlation values shown here actually increase slightly for small decimation factors because decimation serves to partially regularize the overfitted shapes.

**Fig 7.**
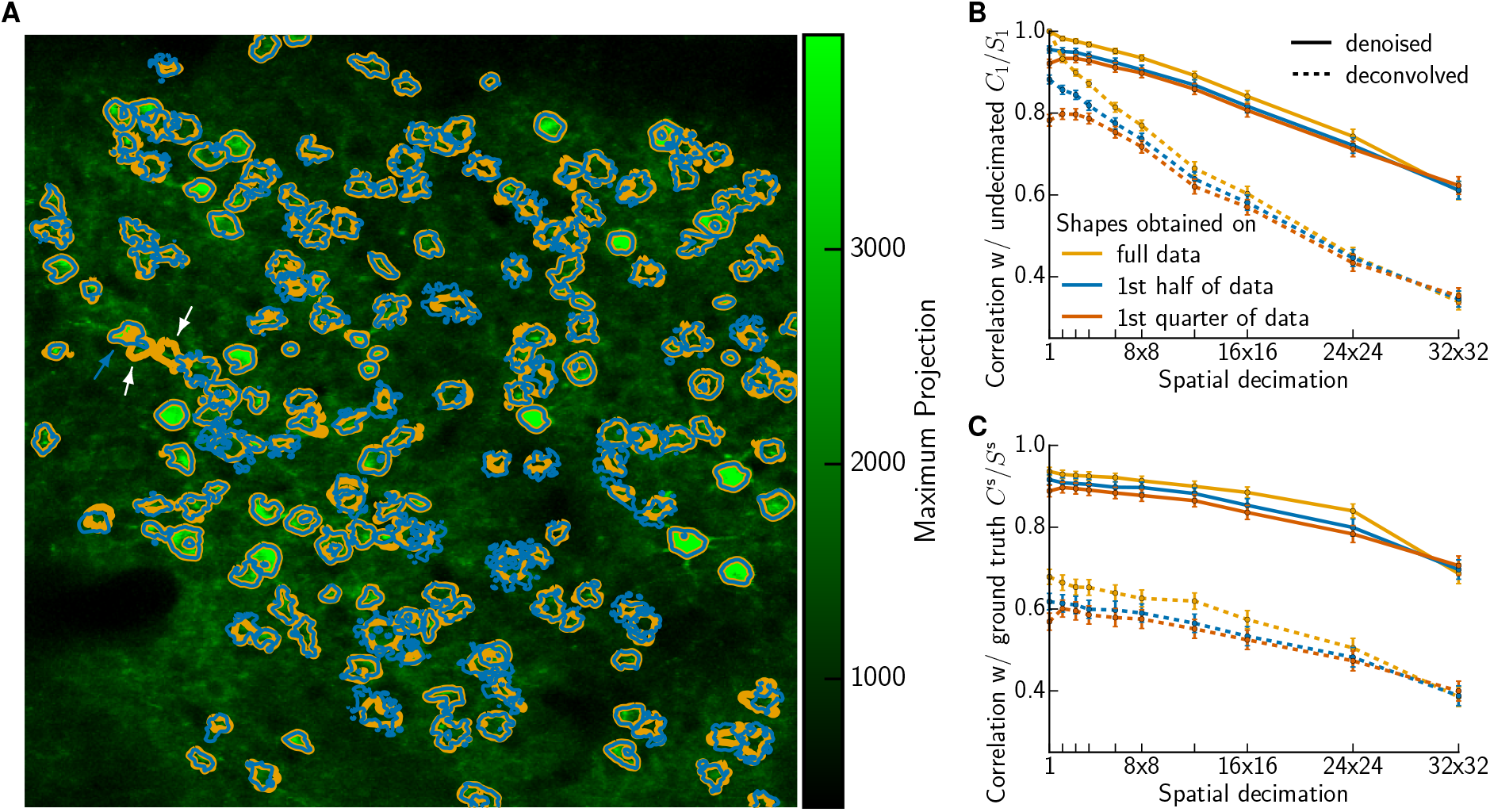
Estimating neural shapes on small initial batch of the data. **(A)** Shapes inferred on the full data (orange) or using only the first half (blue). Two apparently lost suspected ROIs (white arrows) are actually part of another neuron (blue arrow) **(B)** Correlation between traces *C*_*l*_ obtained on decimated data and the reference *C*_*l*_ obtained without any decimation. **(C)** Comparing to simulated ground truth *C*^s^. Traces were obtained on decimated data with reshuffled residuals, otherwise analogous to (B).

Finally, we investigated whether it would be possible to further improve the recovery of *C*_*l*_ from very highly decimated data via an interleaving strategy [20]. Instead of using the same large pixels on each frame, e.g. the ones corresponding to the cyan grid in Fig 8A, we alternate between imaging with the cyan pixelization and the half-shifted orange pixelization. This is helpful, for example, in distinguishing the activity of neurons 20 and 39: based on merely cyan pixels, these neurons contribute to only one and the same pixel and distinguishing their activity would therefore be impossible, whereas the information from the orange pixels afford their separation. It is straightforward to generalize the demixing strategy to exploit the information from the cyan and orange grids at alternating time points (see Methods); Figs. 8B and C show that this interleaving is indeed able to improve recovery at very large decimation levels *l*.

**Fig 8.**
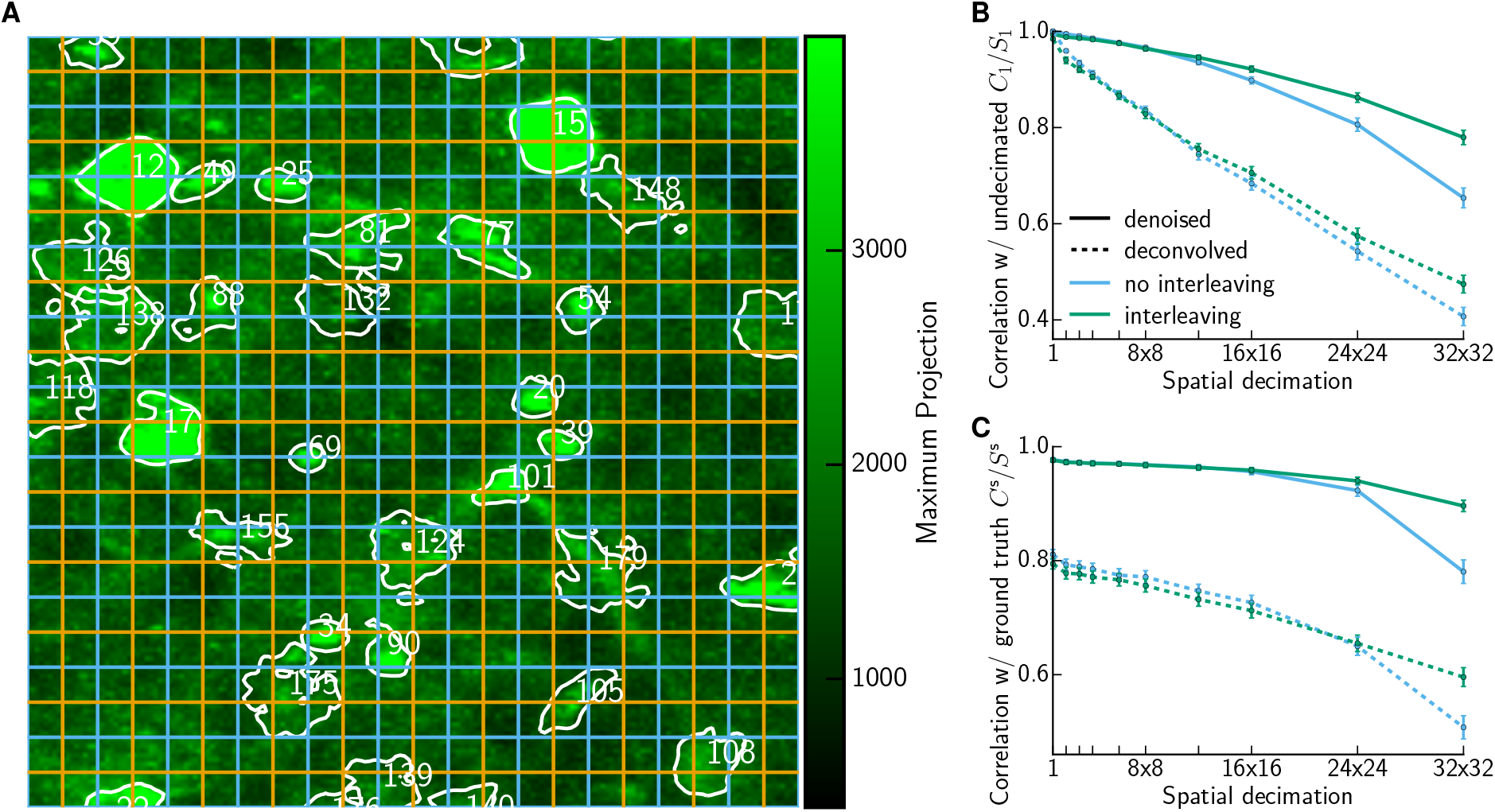
Interleaving improves accuracy of recovered *C*_*l*_ at low spatial resolution. **(A)** Interleaving alternates between pixels corresponding to the cyan and orange grid. **(B)** Correlation between traces *C*_*l*_ obtained on decimated data and the reference *C*_1_ obtained without any decimation. The average correlation (±SEM) decays faster without (cyan) than with interleaving(green). **(C)** Comparing to simulated ground truth *C*^s^. Traces were obtained on decimated data with reshuffled residuals, otherwise analogous to (B).

## Discussion

The basic message of this paper is that standard approaches for imaging calcium responses in large neuronal population — which have historically been optimized so that humans can clearly see cells blink in the resulting video — lead to highly redundant data, and we can exploit this redundancy in several ways. In the first part of the paper, we saw that we can decimate standard calcium imaging video data drastically, to obtain order-of-magnitude speedups in processing time with no loss (and in some cases even some gain) in accuracy of the recovered signals. In the second part of the paper, we saw that, once the cell shapes and locations are identified, we can drastically reduce the spatial resolution of the recording (losing the ability to cleanly identify cells by eye in the resulting heavily-pixelated movies) but still faithfully recover the neural activity of interest. This in turn leads naturally to a proposed two-phase imaging approach (first, identify cell shapes and locations at standard resolution; then image at much lower spatial resolution) that can be seen as an effort to reduce the redundancy of the resulting video data.

We anticipate a number of applications of the results presented here. Regarding the first part of the paper: faster computational processing times are always welcome, of course, but more fundamentally, the faster algorithms developed here open the door towards guided experimental design, in which experimenters can obtain images, process the data quickly, and immediately use this to guide the next experiment. With more effort this closed-loop approach can potentially be implemented in real-time, whether for improving optical brain-machine interfaces [21], or enabling closed-loop optogenetic control of neuronal population dynamics [22,23].

Highly redundant data streams are by definition highly compressible. The results shown in S1 Video and S2 Video illustrate clearly that spatially-decimated image acquisition (the second phase of our two-phase imaging approach) can be seen as a computationally trivial low-loss compression scheme. Again, regarding applications of this compression viewpoint: reductions in memory usage are always welcome — but more fundamentally, this type of compression could for example help enable wireless applications in which bandwidth and power-budget limitations are currently a significant bottleneck [19, 24–26].

Regarding applications of the proposed two-phase imaging approach: we can potentially use this approach to image either more cells, or image cells faster, or some combination of both. In most of the paper we have emphasized the first case, in which we ‘zoom out’ to image larger populations at standard temporal resolution. However, a number of applications require higher temporal resolution. One exciting example is the larval zebrafish, where it is already possible to image the whole brain, but light-sheet whole-brain volumetric imaging rates are low [1] and current efforts are focused on faster acquisition [3,4,27]. Higher temporal resolution is also needed for circuit connectivity inference [28, 29] or the real-time closed-loop applications discussed above, where we need to detect changes in activity as quickly as possible. Finally, genetically encoded voltage indicators [30] may soon enable imaging of neuronal populations with single-cell, millisecond-scale resolution; these indicators are still undergoing intense development [31–34] but when more mature the resulting signals will be much faster than currently-employed calcium indicators and significantly higher temporal resolution will be required to capture these signals.

A number of previous papers can be interpreted in terms of reducing the redundancy of the output image data. Our work can be seen as one example of the general theme of increasing the ratio *N/D*, with *N* denoting the number of imaged neurons and *D* the number of observations per timestep, with demixing algorithms used post hoc to separate the overlapping contributions of each cell to each observed pixel. In a compressed sensing framework, [35] proposed to image randomized projections of the spatial calcium concentration at each timestep, instead of measuring the concentration at individual locations. In [36], [37], and [38], information is integrated primarily across depth, either by creating multiple foci, axially extended point spread functions (PSFs), or both, respectively. In contrast to these methods, [7] instead scanned an enlarged near-isotropic PSF, generated with temporal focusing, to quickly interrogate cells in a single plane at low spatial resolution. This approach is closest in spirit to the one-phase spatially decimated imaging approach analyzed in Figs. 6-8, and could potentially be combined with our two-phase approach to achieve further speed/accuracy gains.

We expect that different strategies for increasing *N/D* will have different advantages in different situations. One advantage of the approach developed here is its apparent simplicity - at least at a conceptual level, we just need to ‘zoom out’ without the need for radically new imaging hardware. Throughout this work we have remained deliberately agnostic regarding the physical implementation of the spatial decimation; all of the decimation results presented here were based on software decimation after acquisition of standard-resolution images. Thus to close we turn now to a discussion of potential experimental caveats.

One critical assumption in our simulations is that the total recorded photon flux per frame is the same for each decimation level *l*. This is a reasonable assumption for light-sheet imaging (assuming we are not limited by laser power or by the peak or average light power on the sample): in this case, increasing the effective pixel size could be achieved easily, either isotropically with a telescope, or anisotropically, with a cylindrical lens or anamorphic prism pair. However, faster whole-brain light-sheet imaging requires faster shifts of the light sheet and imaged focal plane. This challenge can be solved by extended depth-of-field (EDoF) pupil encoding [3,4,27], remote focusing [39], or with an electrically tunable lens [40]. Higher light-sheet imaging rates can also be obtained with swept confocally-aligned planar excitation (SCAPE) microscopy [6]. In short, we believe our proposed two-phase imaging approach fits well with a variety of proven light sheet methods; for similar reasons, the two-phase approach would also fit well with light-field imaging methods [41–43].

In traditional two-photon imaging the situation is more complicated. The image is created by serially sweeping a small, diffraction limited point across the sample. Along the “fast” axis, the beam moves continuously, and the integrated signal across a line is constant, regardless of detection pixelation - the signal is simply partitioned into more or fewer bins. Along the “slow” axis, however, the galvonometers are moved in discrete steps, and low pixel numbers generally mean that portions of the image are not scanned, increasing frame speed, but concomitantly these ‘missed’ areas generate no signal. This consequently reduces the total number of photons collected. Thus to achieve the same photon flux over the larger (lower spatially sampled) pixels, while maintaining the same SNR, we require an enlarged PSF, which maps a larger sampled volume to each pixel. This approach was recently demonstrated to be effective in [7]; alternative strategies for enlarging the PSF could involve fixed diffractive optical elements [44] or spatial light modulator (SLM) systems [45]. Programmable phase-only SLMs offer the additional benefit of being able to dynamically change the size and shape of the excitation PSF, even between frames, which may help disambiguate closely spaced sources, and effectively control the recorded source sparsity.

In any instantiation, maximal imaging speed will be limited by the time required to collect enough photons for adequate SNR, which in turn is limited by photophysics and the light tolerance of the sample. In future work we plan to pursue both light-sheet and 2P implementations of the proposed two-phase imaging approach, to quantify the gains in speed and FOV size that can be realized in practice.

## Methods

### Neural data acquisition

The calcium fluorescence of the whole brain of a larval zebrafish was recorded using light-sheet imaging. It was a transgenic (GCaMP6f) zebrafish embedded in agarose but with the agarose around the tail removed. The fish was in a fictive swimming virtual environment as described in [46]. The closed loop setting, characterized by visual feedback being aligned with the recorded motor activity, was periodically interrupted by open loop phases. Whole-brain activity was recorded for 1,500 seconds with a rate of 2 volumes per second.

In vivo two-photon imaging was performed in a transgenic (GCaMP6s) mouse through a cranial window in visual cortex. The mouse was anesthetized (isoflurane) and head-fixed on a Bruker Ultima in vivo microscope with resonant scanners, and spontaneous activity was recorded. The field of view extended over 350 um × 350 urn and was recorded for 100 seconds with a resolution of 512×512 pixels at 20 frames per second.

### Constrained non-negative matrix factorization

In the case of 2P imaging, the field of view contained *N* = 187 ROIs. It was observed for a total number of *T* = 2,000 timesteps and had a total number of *D* = 512 × 512 pixels. We restricted our analysis of the zebrafish data to a representative patch of size *D* = 96×96 pixels containing *N* = 46 ROIs, extracted from a medial z-layer of the whole-brain light-sheet imaging recording of *T* = 3,000 frames. The observations at any point in time can be vectorized in a single column vector of length *D*; thus all the observations can be described by a *D* × *T* matrix *Y*. Following [8], we model *Y* as

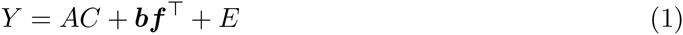

where 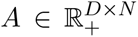 is a spatial matrix that encodes the location and shape of each neuron, 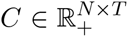 is a temporal matrix that characterizes the calcium concentration of each neuron over time, 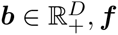, ***f*** ∈ ℝ^*T*^ are nonnegative vectors encoding the background spatial structure and global intensity, respectively, and *E* is additive Gaussian noise with mean zero and diagonal covariance.

For the zebrafish data we ensured that the spatial components are localized, by constraining them to lie within spatial patches (which are not large compared to the size of the cell body) around the neuron centers, thus imposing sparsity on A by construction. Because of the low temporal resolution of these recordings, the inferred neural activity vectors are not expected to be particularly sparse, and therefore we do not impose sparsity in the temporal domain. This leads to the optimization problem

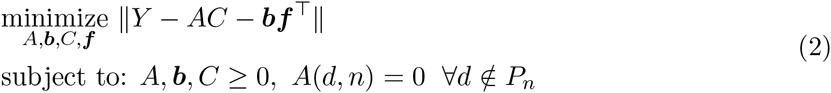

where *P*_*n*_ denotes the *n*-th fixed spatial patch. As discussed in the Results section, we solve this problem by block-coordinate descent, first applied to much smaller decimated data and then using this solution as a warm start for the optimization on the full data. In the resulting Algorithm 1 we appended for conciseness ***b*** and ***f*** as an additional column or row to *A* and *C* respectively.

The matrix products *A*^⊤^*Y* and *CY*^⊤^ in Algorithm 1 are computationally expensive for the full data. These matrix products can also be performed on GPU instead CPU; whereas for the comparatively small 96×96 patches we did not obtain any speed-ups using a GPU, we verified on patches of size 256×256 that some modest overall speedups (a factor of 1.5-2) can be obtained by porting this step to a GPU.

For the decimated data the matrix products are cheap enough to iterate just once over all neurons and instead alternate more often between updating shapes and activities (instead of performing many iterations within HALSactivity or HALSshape in Algorithm 1). In early iterations our estimates of A and C are changing significantly and it is better to perform just one block-coordinate descent step for each neuron to update *A* (and similarly for *C*); for later iterations, and on the full data where it is more expensive to compute *A*^⊤^*Y* and *CY*^⊤^, we increase the inner iterations in HALSactivity or HALSshape.

Algorithm 1 is a constrained version of fast HALS. To further improve on fast HALS, [47] suggested to replace cyclic variable updates with a greedy selection scheme focusing on nonzero elements. This was unnecessary here because most nonzero elements are prespecified by the patches *P*_*n*_; i.e., we are already focusing on the nonzero elements.

Because for the 2P data the observed imaging rate is much higher than the decay rate of the calcium indicator, we constrain the temporal traces *C* to obey the calcium indicator dynamics, to enable further denoising and deconvolution of the data. As in [8], we approximate the calcium concentration dynamics using an autoregressive process of order 2 (AR(2)),

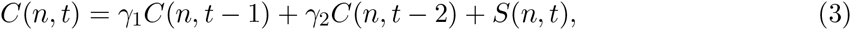

where *S*(*n, t*) is the number of spikes that neuron *n* fired at timestep *t*. This equation can be conveniently expressed in matrix form as *S* = *CG* for a suitable sparse matrix *G*. We estimate the noise level of each pixel *σ*_*d*_ by averaging the power spectral density (PSD) over a range of high frequencies, and estimate the coefficients of the AR(2) process for each cell following [16]. Then we solve for *A*, ***b***, *C*, ***f*** using the following iterative matrix updates:

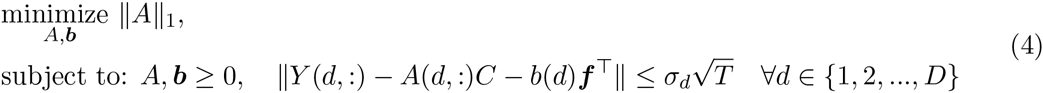

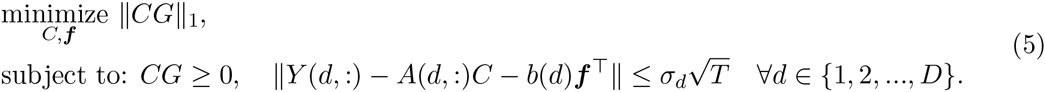

These updates are initialized with the results from constrained fast HALS (Alg 1. They impose sparsity on the spatial as well as temporal components, using the estimates of the noise variance as hard constraints to derive a parameter-free convex program. Following the approach in [48] the spike signal *S* is relaxed from nonnegative integers to arbitrary nonnegative values. The basis pursuit denoising problems in Eqs (4,5) can be solved with one of the methods described in [8]. However, a faster update of the temporal matrix *C* is achieved by using OASIS [16].

### Ranking the obtained components

Every spatial component in *A* was normalized to have unit *ℓ*_2_-norm, with the corresponding temporal component scaled accordingly. Following [8] we then sort the components according to the product of the maximum value that the temporal component attains and the *ℓ*_4_-norm of the corresponding spatial footprints, to penalize overly broad and/or noisy spatial shapes.

### Decimation details

Applying the code of [49] to the raw data we identified the neural shape matrix *A*_1_ and spatial background *b*_1_. We use the convention that the presence of a lower index *l* signifies an estimate and its value the decimation factor, i.e. index *l* = 1 denotes an estimate inferred without decimation. Further, we also obtained the denoised and deconvolved traces *C*_1_, *S*_1_ as well as ***f***_1_. To emulate imaging with lower spatial resolution, spatial decimation was performed by converting *A*_1_ back into a 512 × 512 × *N* tensor (*Y* into a 512 × 512 × *T* video tensor) and calculating the average of non-overlapping patches of size *l* × *l* or *l* × 1 pixels for each of the *N* neural shapes (*T* timesteps). Converting the tensors back to matrices yielded the decimated neural shapes *A*_*l*_ (data *Y*_*l*_). We proceeded analogously for the spatial background to obtain *b*_*l*_. The corresponding temporal traces were estimated by solving Eq (5) (with *Y*_*l*_ replacing *Y, C*_*l*_ replacing *C*, etc.), initializing *C*_*l*_ and ***f***_*l*_ with the result of plain NMF that does not impose temporal constraints, i.e. solving 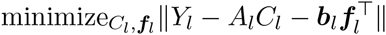 subject to *C*_*l*_ ≥ 0.

In order to obtain the results for 1-phase imaging without previous shape identification we solved Eq (4) for the decimated data *Y*_*l*_, initializing *A*_*l*_, ***b***_*l*_ by decimating *A*_1_, ***b***_1_ and setting the temporal components to *C*_1_, ***f***_1_. With increasing decimation factors an increasing number of shapes got purged and absorbed in the background, reflecting the fact that it would have been difficult to detect all ROIs on low resolution data in the first place. Using the obtained remaining shapes we again solved Eq (5) as above. The correlation values for purged neurons were set to zeros for the mean values reported in Fig 6.

To obtain some form of ground truth (Fig 6D-F, 7C and 8C) we generated a simulated dataset *Y*^s^ by taking the inferred quantities as actual ground truth: *A*^s^:= *A*_1_, ***b***^s^:= ***b***_1_, *C*^s^:= *C*_1_, ***f***^s^:= ***f***_1_. We calculated the residual *Y* − *A*^s^*C*^s^ − ***b***^s^***f***^s⊤^ and reshuffled it independently randomly for each pixel in time. The simulated dataset *Y*^s^ was obtained by adding the reshuffled residual to *A*^s^*C*^s^ + ***b***^s^***f***^s⊤^ and the same analysis as for the original data was performed.

### Interleaving details

Projecting the noise of each pixel unto the neural shapes yields the noise of each neural time series. In practice the latter is estimated based on the noisy trace obtained by projecting the fluorescence data unto the shapes. For interleaved imaging (Fig 8) the shape of each neuron differs between alternating frames due to the varying pixelization. Therefore, instead of using one noise level for all timesteps, we estimated two noise levels *σ*_odd_ and *σ*_even_ based on the PSD for all odd and even frames respectively. The residuals in the noise constraint of the non-negative deconvolution were weighted accordingly by the inverse of the noise level.

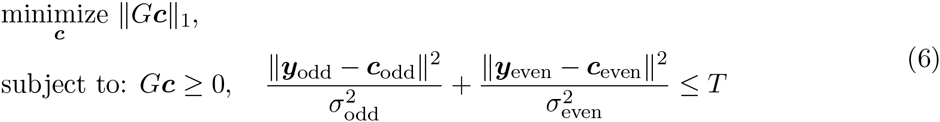

where ***y*** is the noisy fluorescence data of a neuron (cell index suppressed) obtained by subtracting the contribution of all other neurons as well as the background from the spatio-temporal raw pixel fluorescence data and projecting the odd and even frames of the result onto the considered neuron’s shapes ***a***_odd_ and ***a***_even_ respectively. ***y***_odd_ and ***y***_even_ denote the vectors obtained by taking only every second component of ***y*** starting with the first/second respectively. The denoised fluorescence ***c*** is denoted analogously. For simplicity we estimated the coefficients of the AR(2) process based on all frames without separating by noise level.

### Computing environment

All analyses were performed on a MacBook Pro with Intel Core i5-5257U 2.7 GHz CPU and 16 GB RAM. We wrote custom Python scripts that called the Python implementation [49] of CNMF [8]. Our scientific Python installation included Intel Math Kernel Library (MKL) Optimizations for improved performance of vectorized math routines.

## Supporting Information

**S1 Video. Illustration of CNMF and decimation for zebrafish light-sheet imaging data.** The supplementary video shows in the upper row the raw data, its reconstruction based on CNMF, and the residual. It illustrates that all relevant ROIs have been detected and the matrix decomposition afforded by CNMF captures the data well. The lower row shows spatially decimated raw data, corresponding to data acquisition at lower resolution, its reconstruction based on CNMF and knowledge of the neural shapes from an initial cell identification imaging phase, and the residual. It illustrates that demixing paired with cell shape identification captures the data virtually as well as CNMF without decimation, i.e. based on data of low resolution it enables its reconstruction at higher resolution.

**S2 Video. Illustration of CNMF and decimation for mouse 2P imaging data.** All panels of the supplementary video are analogous to S1 Video.

## Acknowledgments

We would like to thank Weiqun Fang for preparing the mouse and Daniel Soudry, Eftychios Pnevmatikakis, Lloyd Russell, Adam Packer, and Jeremy Freeman for helpful conversations. We thank Andrea Giovannucci for his efforts to make a Python implementation of CNMF available.

Part of this work was previously presented at the NIPS (2015) workshop on Statistical Methods for Understanding Neural Systems [50].

Funding for this research was provided by Swiss National Science Foundation (http://www.snf.ch) Research Award P300P2_158428 (JF), the Gruss Lipper Charitable Foundation (http://eglcf.org; DS), Simons Foundation (https://www.simonsfoundation.org) Global Brain Research Awards 325171 (MA, LP), 325398, and 365002 (LP), Howard Hughes Medical Institute (http://hhmi.org; MA), Army Research Office (ARO, https://www.arl.army.mil?page=472) MURI W911NF-12-1-0594 (RY, LP), NIH NIMH R44MH109187 (DSP), NIH BRAIN Initiative (https://www.braininitiative.nih.gov) R01 EB22913 (LP) and R21 EY027592 (DSP, LP), Defense Advanced Research Projects Agency (DARPA, http://www.darpa.mil) N66001-15-C-4032 (SIMPLEX; RY, LP), and a Google Faculty Research award (LP); in addition, this work was supported by the Intelligence Advanced Research Projects Activity (IARPA, https://www.iarpa.gov) via Department of Interior/ Interior Business Center (DoI/IBC) contract numbers D16PC00003 and D16PC00007 (LP). The funders had no role in study design, data collection and analysis, decision to publish, or preparation of the manuscript. The U.S. Government is authorized to reproduce and distribute reprints for Governmental purposes notwithstanding any copyright annotation thereon. Disclaimer: The views and conclusions contained herein are those of the authors and should not be interpreted as necessarily representing the official policies or endorsements, either expressed or implied, of IARPA, DoI/IBC, or the U.S. Government.

